# *Aedes koreicus*, a vector on the rise: pan-European genetic patterns, mitochondrial and draft genome sequencing

**DOI:** 10.1101/2021.12.07.471561

**Authors:** Kornélia Kurucz, Safia Zeghbib, Daniele Arnoldi, Giovanni Marini, Mattia Manica, Alice Michelutti, Fabrizio Montarsi, Isra Deblauwe, Wim Van Bortel, Nathalie Smitz, Wolf Peter Pfitzner, Christina Czajka, Artur Jöst, Katja Kalan, Jana Šušnjar, Vladimir Ivović, Anett Kuczmog, Zsófia Lanszki, Gábor Endre Tóth, Balázs A. Somogyi, Róbert Herczeg, Péter Urbán, Rubén Bueno-Marí, Zoltán Soltész, Gábor Kemenesi

## Abstract

**Background:** The mosquito *Aedes koreicus* (Edwards, 1917) is a recent invader on the European continent that was introduced to several new places since its first detection in 2008. Compared to other exotic *Aedes* mosquitoes with public health significance that invaded Europe during the last decades, this species’ biology, behavior, and dispersal patterns were poorly investigated to date.

**Methodology/Principal Findings:** To understand the species’ population relationships and dispersal patterns within Europe, a fragment of the COI gene was sequenced from 130 mosquitoes, collected from five countries where the species has been introduced and/or established. Oxford Nanopore and Illumina sequencing techniques were combined to generate the first complete nuclear and mitochondrial genomic sequences of *Ae. koreicus* from the European region. The complete genome of *Ae. koreicus* is 879 Mb. COI haplotype analyses identified five major groups (altogether 31 different haplotypes) and revealed a large-scale dispersal pattern between European *Ae. koreicus* populations. Continuous admixture of populations from Belgium, Italy, and Hungary was highlighted, additionally, haplotype diversity and clustering clearly indicate a separation of German sequences from other populations, pointing to an independent introduction of *Ae. koreicus* to Europe. Finally, a genetic expansion signal was identified, suggesting the species might be present in more locations than currently detected.

**Conclusions/Significance:** Our results highlight the importance of genetic research of invasive mosquitoes to understand general dispersal patterns, reveal main dispersal routes and form the baseline of future mitigation actions. The first complete genomic sequence also provides a significant leap in the general understanding of this species, opening the possibility for future genome-related studies, such as the detection of ‘Single Nucleotide Polymorphism’ markers. Considering its public health importance, it is crucial to further investigate the species’ population genetic dynamic, including a larger sampling and additional genomic markers.

**Author Summary:** In the present context of globalization and changing environment, the rapid spread of Invasive Mosquito Species (IMS) across Europe represents a serious public health threat because some species are competent vectors for several pathogens. A better knowledge of the IMS population relationships, demographic trends, and dispersal patterns can help the relevant authorities mitigating further spread. *Aedes koreicus* is an IMS that invaded the continent and has been expanding its geographic range over the last decade. In the present study, one of the most popular DNA marker (COI) was used to investigate the pan-European haplotype diversity and phylogenetic relatedness within and between *Ae. koreicus* populations. Also, the first complete mitochondrial genome and draft nuclear genome of *Ae. koreicus* were generated using combined high-throughput sequencing techniques (Oxford Nanopore, Illumina). This provides a significant leap in the general understanding of this species and opens the possibility for future genomic studies.

## Introduction

The introduction of alien species to new territories and biological invasions represent an increasingly problem in our globalized world. The process has increased with modern trade and travel, which facilitates and intensifies the introduction of invasive alien species (1). Among others, the invasion of mosquitoes of Asian origin has a highlighted impact in Europe (2). Multiple Invasive Mosquito Species (IMS) belonging to the genus *Aedes* have been introduced and have established during the last decades, generating a serious nuisance for people, representing a public health threat due to their potential vector competence for exotic and/or endemic pathogens (3) and a possible but unknown impact on the native ecosystem. Contrary to the Asian tiger mosquito *Aedes albopictus* (Skuse, 1894) and the Asian bush mosquito *Aedes japonicus japonicus* (Theobald, 1901), little is known about another invader, *Aedes koreicus* (Edwards, 1917). This Aedes Invasive Mosquitoes (AIM) has been proposed as an additional global invasive mosquito with good adaptation capability (4). Compared to *Ae. albopictus*, it can tolerate lower temperatures, with faster development and being able to become more abundant than the tiger mosquito during the colder months of the breeding season (5). *Aedes koreicus* is also better adapted to urban environments than the forest-dwelling species *Ae. j. japonicus* (6). These characteristics suggest that *Ae. koreicus* might continue to expand across the continent.

*Aedes koreicus* was first detected out of its native area in Belgium in 2008, in an industrial area in Maasmechelen, province of Limburg, where it is established and overwintering (7). While the population is still present, it does not seem to colonize the surrounding territory (8). Then, the species was found in northeast Italy in 2011, in the province of Belluno, Veneto region (9), where it has rapidly expanded its range. In ten years, the species infested the neighboring provinces i.e. Vicenza, Treviso, and Trento in Trentino-Alto Adige region, and also reached more distant places in the Lombardia, Friuli Venezia Giulia, as well as Liguria regions in northern Italy (10–13). Furthermore, in 2013, the species was introduced to Sochi on the Black Sea coast of Russia (14,15), and observed at the Swiss-Italian border region (16). Meanwhile, a revision of *Ae. japonicus* specimens collected in Slovenia confirmed the introduction of *Ae. koreicus* into the village of Lovrenc na Dravskem Polju, northeast Slovenia in 2013 (17). The species appeared in the federal state of Bavaria, southern Germany in 2015 (18), and a few years later, in Wiesbaden, in the federal state of Hessen, where a population is now established (19–21). At the same time, *Ae. koreicus* was found in Baranya county, southwest Hungary (22), where an overwintering but localized population has established (23), but a few years later (2019), it appeared in the capital city of Budapest, north-central Hungary (24). Recently, *Ae. koreicus* was detected in Western Austria (25), on the Southern Coast of the Crimean Peninsula (26), and in the Republic of Kazakhstan (26).

Since information is restricted to countries where exotic mosquito surveillance activities are implemented, the above mentioned regions only depict our current knowledge of the species distribution (27). Beyond that, knowledge on its dispersal patterns and the basic mechanisms that drive its distribution in Europe is lacking, although, that information is essential for the projection of future scenarios, particularly critical for the implementation of appropriate exotic mosquito monitoring and management plans. Finally, it is still unclear whether the first detected Belgian population represents the source population of further spreading events across Europe or if we faced multiple introduction events from the species’ native range - or a combination of both over time.

In the present study, we analysed correlations between the European populations of *Ae. koreicus* based on population genetic approaches. To do so, an approximately 1000 base pairs fragment of the cytochrome oxidase I (COI) gene was examined at a pan-European scale in order to reveal haplotype diversity and phylogeographic relatedness throughout the known European populations. We also provide the first complete mitochondrial and draft nuclear genome of the species *Ae. koreicus* by long-read sequencing (Oxford Nanopore) coupled with Illumina sequencing methods.

## Methods

### Mosquito samples

To explore the genetic diversity of *Ae. koreicus* within Europe, we analysed a total of 130 mosquito specimens, originating from nine geographically separated populations from five European countries (Belgium, Germany, Hungary, Italy, and Slovenia) where the species is established, or got recently introduced and it is expected to become established soon. The mosquitoes were collected within the framework of local or national monitoring programs, from urban settlements, including parks, gardens, industrial areas, and cemeteries between 2008 and 2020. Eggs were sampled using oviposition traps, larvae originated from water bodies, and adults were collected either with Frommer Updraft Gravid (John W. Hock Comp., Gainesville, FL) or BG-Sentinel traps (Biogents, Regensburg, Germany) in combination with CO_2_ (for more details, see **S1 Table**). The eggs were first hatched in the laboratory, then larvae and adult specimens were preliminarily identified as *Ae. koreicus* using relevant morphological identification keys (2,20).

### DNA extraction and COI sequencing

Nucleic acid from homogenized mosquitoes was extracted individually, using the NucleoSpin DNA Insect Kit (Macherey-Nagel GmbH & Co. KG, Düren, Germany), following the manufacturer’s instructions. The mitochondrial COI region was amplified using the primer combination LCO1490 and UEA8 (targeting a ∼1000 bp long fragment) (26,27), and PCR reactions were performed with the Platinum II Taq Hot-Start DNA Polymerase kit (Invitrogen, CA, USA), using 0.5 µl of the enzyme, 2 µl of 10x buffer, 0.6 µl of 50mM MgCl_2_, 1-1 µl of the forward and reverse primer solutions (10 µM), 0.2 µl dNTP mix (Promega, USA), 14.7 µl nuclease-free water, and used 2 µl of the DNA extract as a template. The PCR protocol included 5 cycles with following parameters: 30 sec at 94 °C, 30 sec at 45 °C, and 1 min at 72 °C, followed by 35 cycles of denaturation for 30 sec at 94 °C, annealing for 1 min at 51 °C, and 1 min of elongation at 72 °C, ending with a final extension step at 72 °C for 10 minutes. PCR products were purified using NEBMonarch PCR and DNA Clean-up Kit (New England Biolabs, USA), following the manufacturer’s instructions. The sequencing was processed by Eurofins Genomics Custom DNA Sequencing service (Eurofins Genomics, Germany).

### Phylogeographic and haplotype analyses

To characterize the dispersal routes within Europe a discrete phylogeographic analysis was implemented using BEAST v1.10.4, under the HKY substitution model, with a strict molecular clock and constant population size parameters. Best substitution model selection was performed using the IQ-TREE web server tool beforehand. To effectively perform a discrete analysis, the Bayesian stochastic search variable selection/BSSVS and the standard symmetric substitution were combined (30,31). The analysis was run for 10.000.000 generations and sampled every 10.000 iterations, 10 % were discarded as burn-in. Thereafter, TRACER v1.6.0 was used to evaluate the Effective Sampling Size (ESS > 200) (32). Moreover, the MCC tree was annotated using TreeAnnotator v1.10.4 and visualized in FigTree v1.4.4. Lastly, SPREAD3 v0.9.7 was used to calculate Bayes Factor (BF) from the resultant log file for each transition rate and to visualize the MCC tree. Thenceforth, DnaSP v6 was used to assess recombination, then to calculate Tajima’s D and generate a haplotype file (33). In the end, a median-joining haplotype network analysis was carried out in POPART software with default setting/epsilon = 0 (34).

### Oxford Nanopore Sequencing

For whole genome sequencing efforts, the DNA of a laboratory-reared female mosquito was used. The mosquito larva for laboratory rearing was collected in June 2019, Pécs, Hungary. Nucleic acid extraction was performed with the NucleoSpin DNA Insect Kit (Macherey-Nagel GmbH & Co. KG, Düren, Germany), following the manufacturer’s instructions. A final concentration of 75.8 ng/µl nucleic acid eluate was obtained and used as a template for the Nanopore library preparation. To this end, the genomic DNA was sheared using g-Tube (Covaris, U.K.) to reach 8 Kbp average fragment size. For the library preparation 1.6 µg sheared genomic DNA was used (Ligation Sequencing Kit, Oxford Nanopore Technologies, Oxford, UK). The DNA was end-prepped with the NEBNext FFPE Repair and Ultra II End Prep Kit and purified using Agencourt AMPure XP (Beckman Coulter Inc., Brea, CA, USA). Then the adapter ligation was performed using NEBNext Quick T4 DNA Ligase. Finally, the adapted library was purified using Agencourt AMPure and the final concentration was determined using the Qubit 3.0 fluorometer. The library was mixed with ONT running buffer and loading beads, and primed on a FLO-MIN106 9.4.1 SpotON Flow Cell attached to the MinION device (running time: 96 hours).

### Illumina Sequencing

A final concentration of 75.8 ng/µl nucleic acid eluate from the same laboratory-reared female mosquito was used for the Illumina sequencing library preparation. The latter was prepared using NEBNext Ultra II FS DNA Library Prep Kit for Illumina (NEB, Ipswitch, MA, USA). Briefly, 400 ng genomic DNA was fragmented, the end was prepped and the adapter was ligated. Then magnetic beads size selection was performed to select 250-300 bp insert size fragments. Finally, the library was amplified according to the manufacturer’s instructions. The quality of the library was checked on a 4200 TapeSation System using D1000 Screen Tape, the quantity was measured on a Qubit 3.0 fluorometer. Illumina sequencing was performed on a NextSeq550 instrument (Illumina, San Diego, CA, USA) with a 2×151 paired-end run configuration.

### Data Filtering

Obtained reads from both high-throughput sequencing techniques were used to create a draft genome assembly. Beforehand, reads were filtered as follow. Porechop (35) with default parameters was used to remove residual ONT adapters. In the case of ONT, reads with mean quality scores > 7 were retained and used for further processing. For the Illumina PE data, adaptor sequences and low-quality reads were filtered out using the program TrimGalore (36). This dataset (BioSample accession: SRR14975286) was normalized with BBnorm (37) to set the coverage to 50x. In total, 32.05 Gb of Nanopore (BioSample accession: SRR14975285) and 99 Gb of raw Illumina PE data were produced.

### Genome Assembly

Nanopore reads were assembled into contigs by Canu (38) with default parameters. We evaluated the completeness of the assembly with the Benchmarking Universal Single-Copy Orthologs (BUSCO, v 4.1.2) (39) software, and we used the Diptera lineage dataset (diptera_odb10), including 56 species and 3285 BUSCOs. The assembly was further polished with the Illumina PE normalized reads. The accuracy of the draft genome was increased with two additional iterations by Pilon (40). These two assemblies were also examined with BUSCO.

## Results and Discussion

### COI haplotype diversity and phylogeography of Ae. koreicus in Europe

In the present paper, we provide the first insight into the dispersal patterns of *Ae. koreicus* within Europe using genetic approaches. Although phylogenetic investigations of the invasive populations of *Ae. albopictus* (41,42) and *Ae. japonicus* (43,44) were undertaken over the last decades, to date, for *Ae. koreicus* only morphological characterizations are available from Belgium (7) and Germany (20). Here we analysed mtDNA COI partial sequences from 130 *Ae. koreicus* mosquitoes that were collected in different countries in Europe (GenBank accessions: OK668709 - OK668837 and JF430393 (**S1 Table**)). A total of 31 different haplotypes were identified, from specimens collected in nine geographically isolated populations of five countries, reflecting high levels of genetic diversity between and within the examined regions. Among them, five haplotypes (Hap_3, Hap_6, Hap_11, Hap_12, Hap_16) are present in higher frequencies, while all the other identified haplotypes refer to single samples (**Fig 1**). The most frequent haplotype (Hap_3, scored in 34 specimens) is also the most widespread. It is present in most of the investigated regions, including Belgium, all the investigated regions of Italy, both Baranya and Budapest in Hungary, as well as in Slovenia. The second most frequent haplotype (Hap_12, scored in 30 specimens) seems geographically restricted to Italy (mainly Trentino-Alto Adige and Veneto regions) and Hungary. Haplotype Hap_6 (scored in 18 specimens) is present mainly in Hungary, as well as in Belgium and Italy. Haplotype Hap_11 (scored in 15 specimens) is restricted to Germany. In Hungary, haplotype Hap_16 was found in 6 sepcimens collected in 2016, 2017, 2019 and 2020. This supports the establishment of an over-wintering population, in accordance with the observational studies (**Fig 1**). The overall haplotype and nucleotide diversity were of 0.849 and 0.005, respectively, and the overall Tajima’s D was of -2,100 (P < 0.001) (for calculation raw data see **Appendix S1**). These divesity values are in accordance with observational data of the species in the continent, altogether suggesting a general population expansion in Europe.

**Fig 1.**
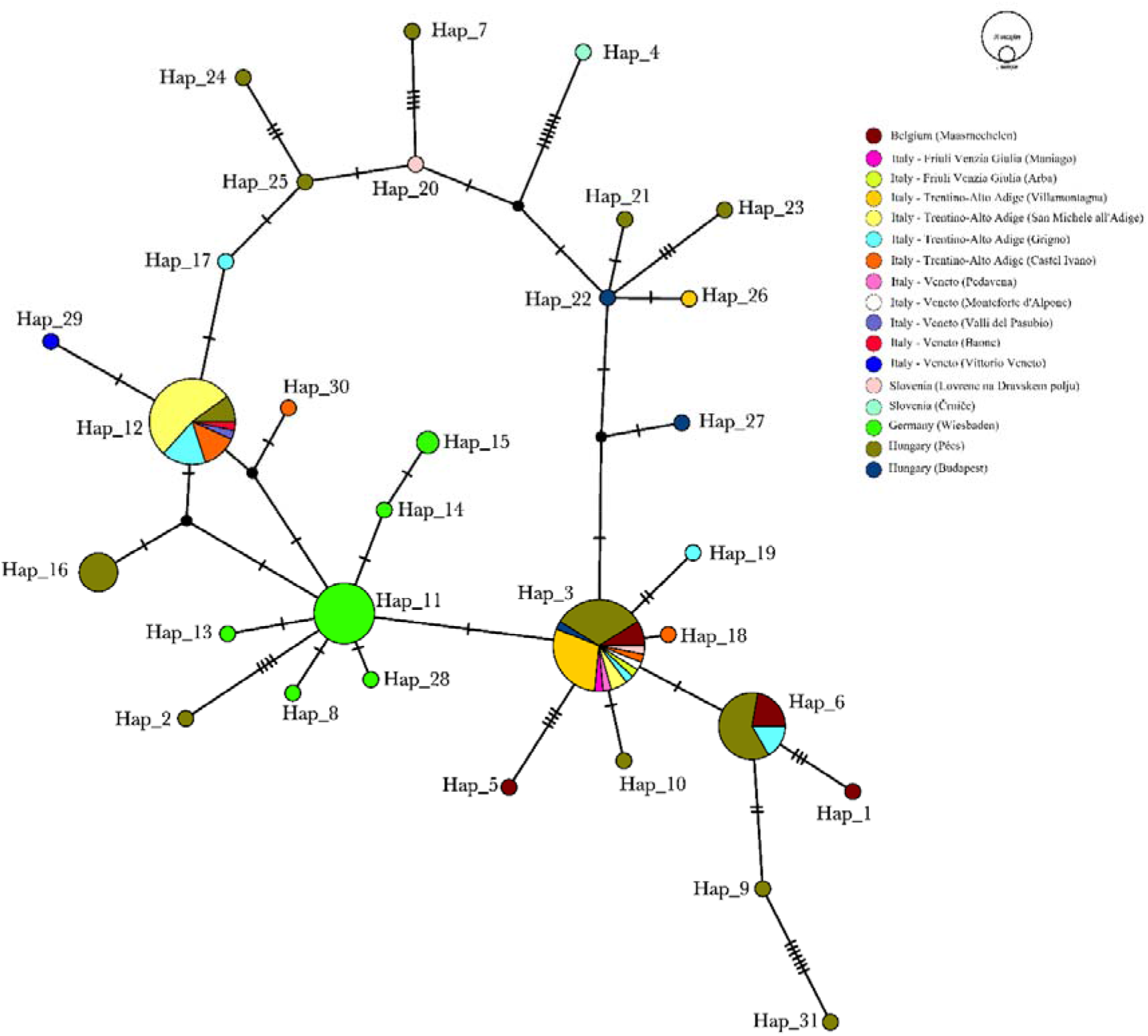
Haplotype network of *Aedes koreicus* constructed with POPART, including 130 mitochondrial COI sequences from specimens collected in five countries (Belgium, Italy, Slovenia, Germany, and Hungary). The 31 haplotypes are indicated on the figure (Hap_1-Hap_31). Different colors represent different sampling sites within the given country.

The phylogeographic analysis clearly indicates the frequent admixture events between European populations and also highlight possible main routes of dispersal (e.g. Belgium-Italy axis) (**Fig 2**). The current Belgian population at Maasmechelen, formerly hypothesized to be the initial introduction site into Europe, consists of a mixture of three haplotypes. Since we do not know if the same haplotypes were already present in 2008, the year of the first observation of *Ae. koreicus*, we cannot draw strong conclusions, but future sequencing efforts, involving more sequences may clarify the genetic diversity and the role of the three identified haplotypes of this study. A previous study on establishing a DNA-barcode reference library for mosquitoes in Belgium also found high intraspecific divergence in *Ae. koreicus*, based on a 658 bp long fragment of the mitochondrial COI gene obtained from two specimens collected in 2008 (45). It can be also possible that Maasmechelen in Belgium was not the initial introduction site for *Ae. koreicus* into Europe, merely it was found there at the earliest, or multiple introductions shaped genetic diversification of the early populations. Future studies are necessary to verify any of these hypotheses.

**Fig 2.**
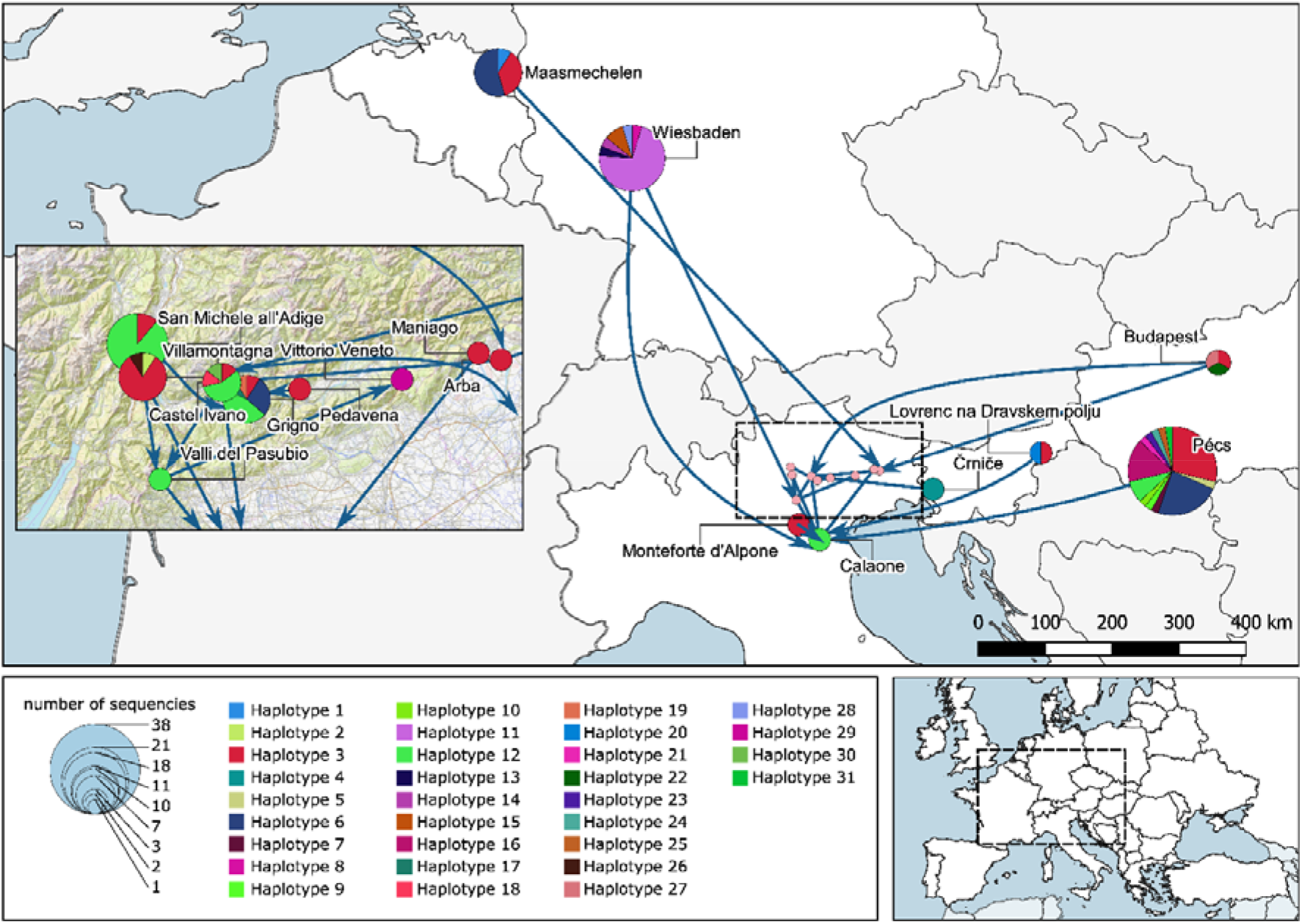
Phylogeography of *Aedes koreicus* based on COI haplotypes in Europe. The geographical placement of pie charts (colored by haplotype distribution) on the map corresponds to the locations where mosquitoes were collected, and the circles size correlates with the number of sequences obtained at each location (for more details about the locations see **S1 Table**). Arrows indicate the potentially most important dispersal routes with significant statistical support (**Appendix S3**).

On the other hand, since the discovery of the invasive *Ae. albopictus* in the north-eastern part of Italy in 1991, several local surveillance and control programs were implemented in the region. Non-colonized (i.e. tiger mosquito-free) areas were also monitored. However, *Ae. koreicus* was first detected in Italy in 2011 (9), suggesting the species was not present before in that specific region. Our analysis indicates that the first population that emerged in Italy probably originated from Belgium. Trading activity of specific goods between Belgium and Italy (back and forth) might be one factor in establishing a major genetic dispersal route between these countries, as presented on **Fig 2**. Admixture was also highlighted between the local populations within Italy, since the analysis revealed four locations where the populations consist of multiple haplotypes (i.e San Michele all’Adige, Villamontagna, Castel Ivano, and Grigno) in the province of Trento (**Fig 2**). Although, these populations are dominantly characterized by the haplotypes Hap_3 and Hap_12 being the most frequent, they share several haplotypes with Belgium and Hungary as well. In the case of Slovenia, the sampled populations share haplotypes with Belgium, Italy, and Hungary, making it hard to define potential routes of introductions, along with the fact that we could involve limited number of samples in our analyses (**Fig 2**).

We revealed the presence of a diverse population in Hungary, which might be explained by common exchange events with the populations in other countries. Since this *Ae. koreicus* population seems to have been established in a short period but is restricted to the city of Pécs (23), the economic center of the Southern Transdanubia region, the high observed haplotype diversity could correlate to the busy international transport of goods through the city. The other Hungarian population at Budapest was represented with a limited sample size in our analyses, but unambiguously conected to the Italian node. The German population displays multiple haplotypes, and the clustering clearly indicates a separation from all other populations and nodes (**Fig 1, Fig 2)**. It may be related to previous observations that the population in Germany was described as the morphological variant of *Ae. koreicus* originating from mainland Korea (20). All other populations in Europe show the morphological characteristics of the Jeju-do Island population (South Korea). The main distinguishing feature between the two variants is that individuals of *Ae. koreicus* from Jeju-do Island have a fifth hind tarsomere with an incomplete pale basal band which is not the case for the specimens from the mainland (47). These perceptions lead to the hypothesis that the German population might be originating from an independent long-distance introduction event.

Based on these results, we hypothesize a silent expansion of the species throughout Europe, with possibly more introduction events than currently identified. Although the routes or directions of the expansion is hard to predict, we pointed to a connectivity between Belgian, Italian and Hungarian populations, and a clear separation of the German population. Considering the yet unverified but hypothesized disease vector potential of *Ae. koreicus* (48– 50), the discovered population interconnectivity underlines the potential rapid dispersal of associated exotic pathogens on these dispersal nodes (51). Meanwhile, the high overall COI diversity may indicate the underrepresentation of the present sample set. Even if we involved specimens from all known established populations in Europe, the present strong expansion signal underlines the importance of extending the sampling efforts for future investigations. Considering the present genetic heterogenity and clustering results, we present similar pathways as a driver of the expansion of *Ae. koreicus* across Europe, as seen in the case of other invasive mosquitoes, e.g. transportation together with specific goods, or with human movements (2,27).

### Complete mitochondrial information and nuclear draft genome sequence

Long-reads obtained with the Oxford Nanopore technology allowed for the construction of the complete mitochondrial and nuclear genomic information, subsequently verified with Illumina short-reads. As a result of combined sequencing methods we were able to retrieve a high-quality mitochondrial genome sequence. Based on the recently published *Ae. koreicus* mitochondrial genome sequenced from a specimen collected from the Korean Peninsula (accession number: MT093832), the final product was annotated. The length of the generated mitogenome (deposited on GenBank with accession number: MZ460582) is 15.843 bp, due to a three basepair long insertion (AAT) at position 9.961. The nucleotide differences between the sequenced Korean specimen (native range) and the European mitochondrial sequence (Hungary, invaded range) are indicated in **Fig 3**.

**Fig 3.**
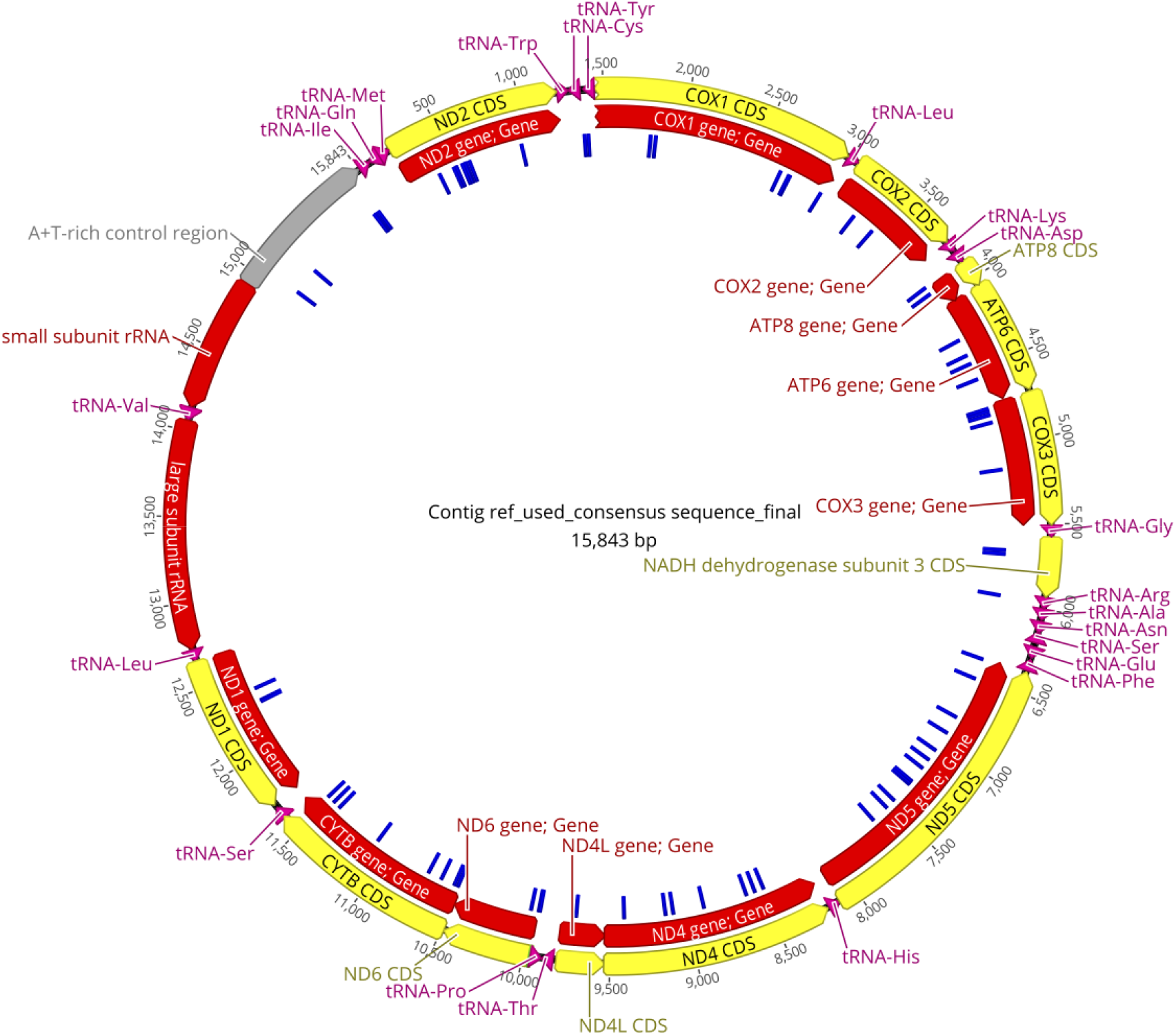
The complete mitochondrial genome of the Hungarian *Aedes koreicus* isolate. Reads generated with Nanopore and Illumina were mapped to the mitochondrial genome with accession number MT093832 with Minimap v.2.17. Gene annotation and visualization were performed in Geneious Prime 2021.1. The yellow circle represents the coding sequence (CDS) or gene product, the red circle represents the sequence of the complete gene, whilst the blue marks in the inner circle represent the single nucleotide differences (n=74) between the genomes of the Hungarian and the Korean *Ae. koreicus* sequences.

Initial assembly of the complete genome yielded a total length of 874 Mb, comprising 65.548 contigs with a contig N50 length of 18 kb. After two iterations, we generated a final genome assembly of 879 Mb (GenBank Submission-ID: 9093984, Sequence Read Archive-IDs: SRR14975285, SRR14975286). BUSCO analysis showed that 2448 (74.52 %) of 3285 Diptera BUSCOs were detected in the final *Ae. koreicus* genome assembly, with 2380 (72.45 %) and 68 (2.07 %) being identified as complete and fragmented respectively (**S2 Table**). The mapping rate test suggested that more than 97.01 % of the normalized Illumina PE reads were properly mapped to the genome, and of these 89.06 % mapped to their mates. These reads covered 98.60 % of the genome assembly. This novel generated draft genome contiguity and N50 lengths lag behind the insect genomes added to public databases in recent years (52). As a neglected invasive mosquito species with yet unverified disease vector potential in Europe, this first draft genome may promote further studies from genome annotation to population genetics.

## Conclusion

Major introduction pathways of exotic mosquitoes and related pathogens are primarily passive and associated with human activities, as global trade and tourism. Since invasions are driven by multiple interacting factors across global, regional, and local scales, we need to investigate the phenomena from a more complex point of view to understand the mechanisms of IMS dispersal. Our study revealed a mixing of European *Ae. koreicus* populations and we also revealed a possible main dispersal route for this species. Interestingly there are two different scenarios in Germany and other European countries, based on the population genetic patterns. At this point we do not know the underlying factors behind the two different situations among these regions, more studies are necessary to get a clearer picture. The first published complete genomic data open the possibility to better understand the genetic features which may facilitate the successful establishment and survival of the species. Based on our results we suggest three key actions of future investigations which we provide the baseline data for. 1. Enforced and harmonized surveillance of this species with common IMS monitoring activities. 2. Re-analysis of previously collected *Ae. japonicus* specimens to reveal possible misidentification. 3. Perform large-scale and more detailed genetic analysis with more samples, using other genetic markers, such as Single Nucleotide Polymorphisms.

## Supporting information Captions

**S1 Table**. Metadata of mosquito samples involved in the present study. A total of 130 COI sequences obtained from Aedes koreicus specimens collected in five European countries between 2008 and 2020 were analyzed. All the sequences are available in GenBank with the referred accession numbers.

**S2 Table**. Quality features of the complete genome sequence of *Aedes koreicus* achieved by Oxford Nanopore and Illumina sequencing. The completeness of the genome assembly was evaluated with the Benchmarking Universal Single-Copy Orthologs (BUSCO, v 4.1.2) software, using the Diptera lineage dataset. C: complete; S: single; D: duplicated; F: fragment; M: missing.

**Appendix S1**. Results of Tajima’s D, Fu and Li’s Neutrality Tests calculations.

**Appendix S2**. Distribution of the 130 investigated *Aedes koreicus* samples among the 31 different haplotypes (Hap#), “Freq” indicates the number of samples belong to the given haplotype, name of the “Sequences” refers to the GenBank Accession Numbers listed in the S1 Table.

**Appendix S3**. Calculated Bayes Factors for specific dispersal routes of Aedes koreicus within the investigated European populations.

## Acknowledgments

The work was done within the framework of the *Aedes* Invasive Mosquitoes COST Action CA17108 (www.aedescost.eu). Mosquito collections in Hungary were supported by the National Research, Development and Innovation Office (NKFIH grant numbers KH-130379, FK-138563, PD-135143, and K-135841), mosquito collections in Slovenia were supported by the Ministry of Health, Ministry of the Environment and Spatial Planning and Slovenian Research Agency (project “Establishment of monitoring of vectors and vector-borne diseases in Slovenia” V3-1903). Mosquito specimens from Veneto and Friuli Venezia (Italy) were collected within the framework of “Invasive mosquito surveillance” as part of activities of the Regional Prevention Plans entitled “Entomological Surveillance of vector-borne diseases” funded by public Health Department of Veneto and Friuli Venezia Giulia Regions. The work in Belgium is part of the MEMO project, funded by the Flemish, Walloon and Brussels regional governments and the Federal Public Service (FPS) Public Health, Food Chain Safety and Environment in the context of the National Environment and Health Action Plan (NEHAP) (Belgium) (tender number CES-2016-02). The Outbreak Research Team of the Institute of Tropical Medicine is financially supported by the Department of Economy, Science and Innovation of the Flemish government. K.K. and G.K. were supported by the Janos Bolyai Research Scholarship of the Hungarian Academy of Sciences and by the ÚNKP-19-4-PTE-264, ÚNKP-20-5-PTE-597, and the ÚNKP-21-5-PTE-1350 New National Excellence Program of the Ministry for Innovation and Technology. S.Z, G.E.T., and Z.L. were supported by the Biological and Sportbiological Doctoral School of the University of Pécs, Hungary. The Barcoding Facility for Organisms and Tissues of Policy Concern (BopCo: http://bopco.myspecies.info/) is financed by the Belgian Science Policy Office (BELSPO) as Belgian federal in-kind contribution to the European Research Infrastructure Consortium “LifeWatch”. R. H. was supported by the grants GINOP-2.3.4-15-2020-00010, GINOP-2.3.1-20-2020-00001 and Educating Experts of the Future: Developing Bioinformatics and Biostatistics competencies of European Biomedical Students (BECOMING, 2019–1-HU01-KA203–061251). Bioinformatics infrastructure was supported by ELIXIR Hungary (http://elixir-hungary.org/). Next generation sequencing and bioinformatics data analyses were partially performed by the Genomics and Bioinformatics Core Faciity of the Szentágothai Research Centre at the University of Pécs. The authors would like to thank the local technicians for their support in mosquito collection management.

## Author Contributions

**Conceptualization:** Kornélia Kurucz, Gábor Kemenesi.

**Data curation:** Kornélia Kurucz, Safia Zeghbib, Róbert Herczeg, Gábor Endre Tóth.

**Formal analysis**: Safia Zeghbib, Gábor Endre Tóth, Róbert Herczeg.

**Funding acquisition:** Kornélia Kurucz, Gábor Kemenesi.

**Investigation:** Kornélia Kurucz, Daniele Arnoldi, Giovanni Marini, Mattia Manica, Alice Michelutti, Fabrizio Montarsi, Isra Deblauwe, Wim Van Bortel, Nathalie Smitz, Wolf Peter Pfitzner, Christina Czajka, Artur Jöst, Katja Kalan, Jana Šušnjar, Vladimir Ivović, Anett Kuczmog, Zsófia Lanszki, Rubén Bueno-Marí, Péter Urbán, Zoltán Soltész, Gábor Kemenesi.

**Methodology:** Gábor Kemenesi, Safia Zeghbib, Gábor Endre Tóth.

**Project administration:** Kornélia Kurucz.

**Resources:** Kornélia Kurucz, Daniele Arnoldi, Giovanni Marini, Mattia Manica, Alice Michelutti, Fabrizio Montarsi, Isra Deblauwe, Wim Van Bortel, Nathalie Smitz, Wolf Peter Pfitzner, Christina Czajka, Artur Jöst, Katja Kalan, Jana Šušnjar, Vladimir Ivović, Zoltán Soltész, Gábor Kemenesi.

**Supervision:** Kornélia Kurucz, Gábor Kemenesi.

**Validation:** Gábor Kemenesi

**Visualization:** Gábor Endre Tóth, Balázs A. Somogyi

**Writing– original draft:** Kornélia Kurucz, Gábor Kemenesi

**Writing– review & editing:** Kornélia Kurucz, Safia Zeghbib, Daniele Arnoldi, Giovanni Marini, Mattia Manica, Alice Michelutti, Fabrizio Montarsi, Isra Deblauwe, Wim Van Bortel, Nathalie Smitz, Wolf Peter Pfitzner, Christina Czajka, Artur Jöst, Katja Kalan, Jana Šušnjar, Vladimir Ivović, Anett Kuczmog, Zsófia Lanszki, Gábor Endre Tóth, Balázs A. Somogyi, Róbert Herczeg, Péter Urbán, Rubén Bueno-Marí, Zoltán Soltész, Gábor Kemenesi.

