## Supporting information for "*Aedes koreicus*, a vector on the rise: pan-European genetic patterns, mitochondrial and draft genome sequencing"

**S1 Table.** Metadata of mosquito samples involved in the present study. A total of 130 COI sequences obtained from *Aedes koreicus* specimens collected in five European countries between 2008 and 2020 were analyzed. All the sequences are available in GenBank with the referred accession numbers.

| **Sample ID** | **COI GenBank Accession No** | **Country** | **Region** | **Province** | **Collection Place** | **Habitat** | **Coordinates** | | **Sample Collection Date** |
| --- | --- | --- | --- | --- | --- | --- | --- | --- | --- |
| HU_Bar_2016_130 | OK668713 | Hungary | Southern Transdanubia | Baranya | Pécs | urban | 46.139391 | 18.224344 | 2016.08.03. |
| HU_Bar_2016_162 | OK668736 | Hungary | Southern Transdanubia | Baranya | Pécs | urban | 46.139391 | 18.224344 | 2016.08.24. |
| HU_Bar_2016_187 | OK668767 | Hungary | Southern Transdanubia | Baranya | Pécs | urban | 46.139391 | 18.224344 | 2016.09.14. |
| HU_Bar_2017_214 | OK668715 | Hungary | Southern Transdanubia | Baranya | Pécs | urban | 46.139391 | 18.224344 | 2017.05.31. |
| HU_Bar_2017_227 | OK668714 | Hungary | Southern Transdanubia | Baranya | Pécs | urban | 46.139391 | 18.224344 | 2017.06.07. |
| HU_Bar_2017_241 | OK668730 | Hungary | Southern Transdanubia | Baranya | Pécs | urban | 46.139391 | 18.224344 | 2017.06.14. |
| HU_Bar_2017_354 | OK668737 | Hungary | Southern Transdanubia | Baranya | Pécs | urban | 46.139391 | 18.224344 | 2017.08.09. |
| HU_Bar_2017_371 | OK668816 | Hungary | Southern Transdanubia | Baranya | Pécs | urban | 46.139391 | 18.224344 | 2017.08.16. |
| HU_Bar_2017_374 | OK668748 | Hungary | Southern Transdanubia | Baranya | Pécs | urban | 46.139391 | 18.224344 | 2017.08.16. |
| HU_Bar_2017_378 | OK668764 | Hungary | Southern Transdanubia | Baranya | Pécs | urban | 46.139391 | 18.224344 | 2017.08.16. |
| HU_Bar_2017_397 | OK668778 | Hungary | Southern Transdanubia | Baranya | Pécs | urban | 46.139391 | 18.224344 | 2017.08.30. |
| HU_Bar_2018_430 | OK668834 | Hungary | Southern Transdanubia | Baranya | Pécs | urban | 46.139391 | 18.224344 | 2018.05.16. |
| HU_Bar_2018_466 | OK668747 | Hungary | Southern Transdanubia | Baranya | Pécs | urban | 46.139391 | 18.224344 | 2018.05.30. |
| HU_Bar_2018_596 | OK668712 | Hungary | Southern Transdanubia | Baranya | Pécs | urban | 46.139391 | 18.224344 | 2018.06.27. |
| HU_Bar_2018_626 | OK668831 | Hungary | Southern Transdanubia | Baranya | Pécs | urban | 46.139391 | 18.224344 | 2018.07.04. |
| HU_Bar_2018_660 | OK668828 | Hungary | Southern Transdanubia | Baranya | Pécs | urban | 46.139391 | 18.224344 | 2018.07.11. |
| HU_Bar_2018_684 | OK668743 | Hungary | Southern Transdanubia | Baranya | Pécs | urban | 46.139391 | 18.224344 | 2018.08.01. |
| HU_Bar_2018_697 | OK668819 | Hungary | Southern Transdanubia | Baranya | Pécs | urban | 46.139391 | 18.224344 | 2018.08.15. |
| HU_Bar_2018_699 | OK668833 | Hungary | Southern Transdanubia | Baranya | Pécs | urban | 46.139391 | 18.224344 | 2018.08.22. |
| HU_Bar_2018_708 | OK668709 | Hungary | Southern Transdanubia | Baranya | Pécs | urban | 46.139391 | 18.224344 | 2018.08.29. |
| HU_Bar_2018_715 | OK668753 | Hungary | Southern Transdanubia | Baranya | Pécs | urban | 46.139391 | 18.224344 | 2018.09.05. |
| HU_Bar_2019_740 | OK668750 | Hungary | Southern Transdanubia | Baranya | Pécs | urban | 46.139391 | 18.224344 | 2019.05.15. |
| HU_Bar_2019_750 | OK668751 | Hungary | Southern Transdanubia | Baranya | Pécs | urban | 46.139391 | 18.224344 | 2019.05.15. |
| HU_Bar_2019_753 | OK668795 | Hungary | Southern Transdanubia | Baranya | Pécs | urban | 46.139391 | 18.224344 | 2019.05.15. |
| HU_Bar_2019_788 | OK668822 | Hungary | Southern Transdanubia | Baranya | Pécs | urban | 46.139391 | 18.224344 | 2019.06.05. |
| HU_Bar_2019_826 | OK668821 | Hungary | Southern Transdanubia | Baranya | Pécs | urban | 46.139391 | 18.224344 | 2019.06.12. |
| HU_Bar_2019_912 | OK668749 | Hungary | Southern Transdanubia | Baranya | Pécs | urban | 46.139391 | 18.224344 | 2019.07.03. |
| HU_Bar_2019_978 | OK668744 | Hungary | Southern Transdanubia | Baranya | Pécs | urban | 46.139391 | 18.224344 | 2019.07.17. |
| HU_Bar_2019_1030 | OK668745 | Hungary | Southern Transdanubia | Baranya | Pécs | urban | 46.139391 | 18.224344 | 2019.07.17. |
| HU_Bar_2019_1072 | OK668742 | Hungary | Southern Transdanubia | Baranya | Pécs | urban | 46.139391 | 18.224344 | 2019.08.22. |
| HU_Bar_2019_1079 | OK668777 | Hungary | Southern Transdanubia | Baranya | Pécs | urban | 46.139391 | 18.224344 | 2019.08.28. |
| HU_Bar_2019_1080 | OK668746 | Hungary | Southern Transdanubia | Baranya | Pécs | urban | 46.139391 | 18.224344 | 2019.08.28. |
| HU_Bar_2019_1082 | OK668738 | Hungary | Southern Transdanubia | Baranya | Pécs | urban | 46.139391 | 18.224344 | 2019.09.04. |
| HU_Bar_2019_1085 | OK668716 | Hungary | Southern Transdanubia | Baranya | Pécs | urban | 46.139391 | 18.224344 | 2019.09.04. |
| HU_Bar_2019_1088 | OK668766 | Hungary | Southern Transdanubia | Baranya | Pécs | urban | 46.139391 | 18.224344 | 2019.09.11. |
| HU_Bar_2019_1090 | OK668765 | Hungary | Southern Transdanubia | Baranya | Pécs | urban | 46.139391 | 18.224344 | 2019.09.11. |
| HU_Bar_2020_2 | OK668717 | Hungary | Southern Transdanubia | Baranya | Pécs | urban | 46.139391 | 18.224344 | 2020.08.27. |
| HU_Bar_2020_3 | OK668763 | Hungary | Southern Transdanubia | Baranya | Pécs | urban | 46.139391 | 18.224344 | 2020.09.11. |
| HU_Bud_2020_1 | OK668752 | Hungary | Central-Hungary | Pest | Budapest | urban | 47.519286 | 19.077579 | 2020.09.29. |
| HU_Bud_2020_2 | OK668710 | Hungary | Central-Hungary | Pest | Budapest | urban | 47.519286 | 19.077579 | 2020.09.29. |
| HU_Bud_2020_4 | OK668711 | Hungary | Central-Hungary | Pest | Budapest | urban | 47.519286 | 19.077579 | 2020.09.29. |
| SN_Dra_2013_539 | OK668820 | Slovenia | Drava |  | Lovrenc na Dravskem polju | urban | 46.373849 | 15.783902 | 2013.09.01. |
| SN_Dra_2013_540 | OK668759 | Slovenia | Drava |  | Lovrenc na Dravskem polju | urban | 46.373849 | 15.783902 | 2013.09.01. |
| SN_Vip_2019_01 | OK668835 | Slovenia | Goriška | Vipava valley | Črniče | cemetery | 45.905009 | 13.777402 | 2019.09.04. |
| IT_Ven_2020_1 | OK668785 | Italy | Veneto | Padova | Baone | urban | 45.249579 | 11.673661 | 2019.09.27. |
| IT_Ven_2020_2 | OK668720 | Italy | Veneto | Belluno | Pedavena | urban | 46.051107 | 11.872342 | 2020.05.14. |
| IT_Ven_2020_3 | OK668779 | Italy | Veneto | Vicenza | Valli del Pasubio | urban | 45.765623 | 11.238465 | 2020.05.14. |
| IT_Ven_2020_4 | OK668829 | Italy | Veneto | Treviso | Vittorio Veneto | urban | 46.081524 | 12.336247 | 2020.06.07. |
| IT_Fri_2020_5 | OK668719 | Italy | Friuli Venzia Giulia | Pordenone | Maniago | urban | 46.164100 | 12.683844 | 2020.06.19. |
| IT_Fri_2020_6 | OK668727 | Italy | Friuli Venzia Giulia | Pordenone | Arba | urban | 46.143553 | 12.785998 | 2020.06.19. |
| IT_Ven_2020_8 | OK668731 | Italy | Veneto | Verona | Monteforte d'Alpone | urban | 45.444958 | 11.285270 | 2020.06.30. |
| IT_Tre_2019_2 | OK668790 | Italy | Trentino-Alto Adige | Trento | Grigno | urban | 46.019535 | 11.631182 | 2019.10.05. |
| IT_Tre_2019_4 | OK668789 | Italy | Trentino-Alto Adige | Trento | Grigno | urban | 46.019535 | 11.631182 | 2019.10.05. |
| IT_Tre_2019_5 | OK668784 | Italy | Trentino-Alto Adige | Trento | Grigno | urban | 46.019535 | 11.631182 | 2019.10.05. |
| IT_Tre_2019_6 | OK668758 | Italy | Trentino-Alto Adige | Trento | Grigno | urban | 46.019535 | 11.631182 | 2019.10.05. |
| IT_Tre_2019_8 | OK668739 | Italy | Trentino-Alto Adige | Trento | Grigno | urban | 46.019535 | 11.631182 | 2019.10.05. |
| IT_Tre_2019_11 | OK668794 | Italy | Trentino-Alto Adige | Trento | Grigno | urban | 46.019535 | 11.631182 | 2019.10.05. |
| IT_Tre_2019_14 | OK668741 | Italy | Trentino-Alto Adige | Trento | Grigno | urban | 46.019535 | 11.631182 | 2019.10.05. |
| IT_Tre_2019_16 | OK668740 | Italy | Trentino-Alto Adige | Trento | Grigno | urban | 46.019535 | 11.631182 | 2019.10.05. |
| IT_Tre_2019_18 | OK668787 | Italy | Trentino-Alto Adige | Trento | Grigno | urban | 46.019535 | 11.631182 | 2019.10.05. |
| IT_Tre_2019_19 | OK668788 | Italy | Trentino-Alto Adige | Trento | Grigno | urban | 46.019535 | 11.631182 | 2019.10.05. |
| IT_Tre_2019_20 | OK668726 | Italy | Trentino-Alto Adige | Trento | Grigno | urban | 46.019535 | 11.631182 | 2019.10.05. |
| IT_Tre_2020_21 | OK668826 | Italy | Trentino-Alto Adige | Trento | Castel Ivano | urban | 46.072517 | 11.518469 | 2020.05.26. |
| IT_Tre_2020_22 | OK668757 | Italy | Trentino-Alto Adige | Trento | Castel Ivano | urban | 46.072517 | 11.518469 | 2020.05.26. |
| IT_Tre_2020_26 | OK668800 | Italy | Trentino-Alto Adige | Trento | Castel Ivano | urban | 46.072517 | 11.518469 | 2020.05.26. |
| IT_Tre_2020_29 | OK668825 | Italy | Trentino-Alto Adige | Trento | Castel Ivano | urban | 46.072517 | 11.518469 | 2020.05.26. |
| IT_Tre_2020_30 | OK668786 | Italy | Trentino-Alto Adige | Trento | Castel Ivano | urban | 46.072517 | 11.518469 | 2020.05.26. |
| IT_Tre_2020_32 | OK668791 | Italy | Trentino-Alto Adige | Trento | Castel Ivano | urban | 46.072517 | 11.518469 | 2020.05.26. |
| IT_Tre_2020_35 | OK668796 | Italy | Trentino-Alto Adige | Trento | Castel Ivano | urban | 46.072517 | 11.518469 | 2020.05.26. |
| IT_Tre_2020_41 | OK668725 | Italy | Trentino-Alto Adige | Trento | San Michele all'Adige | urban | 46.193126 | 11.135434 | 2020.05.26. |
| IT_Tre_2020_44 | OK668760 | Italy | Trentino-Alto Adige | Trento | San Michele all'Adige | urban | 46.193126 | 11.135434 | 2020.05.26. |
| IT_Tre_2020_45 | OK668815 | Italy | Trentino-Alto Adige | Trento | San Michele all'Adige | urban | 46.193126 | 11.135434 | 2020.05.26. |
| IT_Tre_2020_52 | OK668781 | Italy | Trentino-Alto Adige | Trento | San Michele all'Adige | urban | 46.193126 | 11.135434 | 2020.05.26. |
| IT_Tre_2020_53 | OK668783 | Italy | Trentino-Alto Adige | Trento | San Michele all'Adige | urban | 46.193126 | 11.135434 | 2020.05.26. |
| IT_Tre_2020_54 | OK668793 | Italy | Trentino-Alto Adige | Trento | San Michele all'Adige | urban | 46.193126 | 11.135434 | 2020.05.26. |
| IT_Tre_2020_55 | OK668797 | Italy | Trentino-Alto Adige | Trento | San Michele all'Adige | urban | 46.193126 | 11.135434 | 2020.05.26. |
| IT_Tre_2020_56 | OK668801 | Italy | Trentino-Alto Adige | Trento | San Michele all'Adige | urban | 46.193126 | 11.135434 | 2020.05.26. |
| IT_Tre_2020_57 | OK668782 | Italy | Trentino-Alto Adige | Trento | San Michele all'Adige | urban | 46.193126 | 11.135434 | 2020.05.26. |
| IT_Tre_2020_58 | OK668780 | Italy | Trentino-Alto Adige | Trento | San Michele all'Adige | urban | 46.193126 | 11.135434 | 2020.05.26. |
| IT_Tre_2020_59 | OK668798 | Italy | Trentino-Alto Adige | Trento | San Michele all'Adige | urban | 46.193126 | 11.135434 | 2020.05.26. |
| IT_Tre_2020_60 | OK668802 | Italy | Trentino-Alto Adige | Trento | San Michele all'Adige | urban | 46.193126 | 11.135434 | 2020.05.26. |
| IT_Tre_2020_61 | OK668827 | Italy | Trentino-Alto Adige | Trento | San Michele all'Adige | urban | 46.193126 | 11.135434 | 2020.05.26. |
| IT_Tre_2020_62 | OK668803 | Italy | Trentino-Alto Adige | Trento | San Michele all'Adige | urban | 46.193126 | 11.135434 | 2020.05.26. |
| IT_Tre_2020_63 | OK668804 | Italy | Trentino-Alto Adige | Trento | San Michele all'Adige | urban | 46.193126 | 11.135434 | 2020.05.26. |
| IT_Tre_2020_64 | OK668814 | Italy | Trentino-Alto Adige | Trento | San Michele all'Adige | urban | 46.193126 | 11.135434 | 2020.05.26. |
| IT_Tre_2020_65 | OK668799 | Italy | Trentino-Alto Adige | Trento | San Michele all'Adige | urban | 46.193126 | 11.135434 | 2020.05.26. |
| IT_Tre_2020_66 | OK668792 | Italy | Trentino-Alto Adige | Trento | San Michele all'Adige | urban | 46.193126 | 11.135434 | 2020.05.26. |
| IT_Tre_2020_68 | OK668761 | Italy | Trentino-Alto Adige | Trento | Villamontagna | urban | 46.089603 | 11.159079 | 2020.05.26. |
| IT_Tre_2020_71 | OK668754 | Italy | Trentino-Alto Adige | Trento | Villamontagna | urban | 46.089603 | 11.159079 | 2020.05.26. |
| IT_Tre_2020_72 | OK668755 | Italy | Trentino-Alto Adige | Trento | Villamontagna | urban | 46.089603 | 11.159079 | 2020.05.26. |
| IT_Tre_2020_73 | OK668824 | Italy | Trentino-Alto Adige | Trento | Villamontagna | urban | 46.089603 | 11.159079 | 2020.05.26. |
| IT_Tre_2020_74 | OK668823 | Italy | Trentino-Alto Adige | Trento | Villamontagna | urban | 46.089603 | 11.159079 | 2020.05.26. |
| IT_Tre_2020_75 | OK668836 | Italy | Trentino-Alto Adige | Trento | Villamontagna | urban | 46.089603 | 11.159079 | 2020.05.26. |
| IT_Tre_2020_76 | OK668837 | Italy | Trentino-Alto Adige | Trento | Villamontagna | urban | 46.089603 | 11.159079 | 2020.05.26. |
| IT_Tre_2020_77 | OK668724 | Italy | Trentino-Alto Adige | Trento | Villamontagna | urban | 46.089603 | 11.159079 | 2020.05.26. |
| IT_Tre_2020_78 | OK668729 | Italy | Trentino-Alto Adige | Trento | Villamontagna | urban | 46.089603 | 11.159079 | 2020.05.26. |
| IT_Tre_2020_79 | OK668718 | Italy | Trentino-Alto Adige | Trento | Villamontagna | urban | 46.089603 | 11.159079 | 2020.05.26. |
| IT_Tre_2020_81 | OK668728 | Italy | Trentino-Alto Adige | Trento | Villamontagna | urban | 46.089603 | 11.159079 | 2020.05.26. |
| DE_Hes_2019_1 | OK668772 | Germany | Hessen | Wiesbaden | Wiesbaden | cemetery | 50.05853 | 8.268892 | 2019.08.08. |
| DE_Hes_2019_2 | OK668776 | Germany | Hessen | Wiesbaden | Wiesbaden | cemetery | 50.05853 | 8.268892 | 2019.08.08. |
| DE_Hes_2019_3 | OK668805 | Germany | Hessen | Wiesbaden | Wiesbaden | cemetery | 50.05853 | 8.268892 | 2019.08.29. |
| DE_Hes_2019_4 | OK668775 | Germany | Hessen | Wiesbaden | Wiesbaden | cemetery | 50.05853 | 8.268892 | 2019.08.29. |
| DE_Hes_2019_5 | OK668773 | Germany | Hessen | Wiesbaden | Wiesbaden | cemetery | 50.05853 | 8.268892 | 2019.08.29. |
| DE_Hes_2019_6 | OK668769 | Germany | Hessen | Wiesbaden | Wiesbaden | cemetery | 50.05853 | 8.268892 | 2019.08.29. |
| DE_Hes_2019_7 | OK668770 | Germany | Hessen | Wiesbaden | Wiesbaden | cemetery | 50.05853 | 8.268892 | 2019.08.29. |
| DE_Hes_2019_8 | OK668830 | Germany | Hessen | Wiesbaden | Wiesbaden | cemetery | 50.05853 | 8.268892 | 2019.08.29. |
| DE_Hes_2019_9 | OK668806 | Germany | Hessen | Wiesbaden | Wiesbaden | cemetery | 50.05853 | 8.268892 | 2019.08.29. |
| DE_Hes_2019_10 | OK668771 | Germany | Hessen | Wiesbaden | Wiesbaden | cemetery | 50.05853 | 8.268892 | 2019.08.29. |
| DE_Hes_2019_11 | OK668774 | Germany | Hessen | Wiesbaden | Wiesbaden | cemetery | 50.05853 | 8.268892 | 2019.08.29. |
| DE_Hes_2019_12 | OK668768 | Germany | Hessen | Wiesbaden | Wiesbaden | cemetery | 50.05853 | 8.268892 | 2019.08.29. |
| DE_Hes_2019_13 | OK668817 | Germany | Hessen | Wiesbaden | Wiesbaden | cemetery | 50.015663 | 8.281782 | 2019.08.15. |
| DE_Hes_2019_26 | OK668818 | Germany | Hessen | Wiesbaden | Wiesbaden | cemetery | 50.097884 | 8.270688 | 2019.09.27. |
| DE_Hes_2019_27 | OK668808 | Germany | Hessen | Wiesbaden | Wiesbaden | cemetery | 50.097884 | 8.270688 | 2019.09.27. |
| DE_Hes_2019_28 | OK668807 | Germany | Hessen | Wiesbaden | Wiesbaden | cemetery | 50.097884 | 8.270688 | 2019.09.27. |
| DE_Hes_2019_29 | OK668812 | Germany | Hessen | Wiesbaden | Wiesbaden | cemetery | 50.097884 | 8.270688 | 2019.09.27. |
| DE_Hes_2019_30 | OK668810 | Germany | Hessen | Wiesbaden | Wiesbaden | cemetery | 50.097884 | 8.270688 | 2019.09.27. |
| DE_Hes_2019_31 | OK668811 | Germany | Hessen | Wiesbaden | Wiesbaden | cemetery | 50.097884 | 8.270688 | 2019.09.27. |
| DE_Hes_2019_32 | OK668809 | Germany | Hessen | Wiesbaden | Wiesbaden | cemetery | 50.076252 | 8.181752 | 2019.08.29. |
| DE_Hes_2019_33 | OK668813 | Germany | Hessen | Wiesbaden | Wiesbaden | cemetery | 50.076252 | 8.181752 | 2019.08.29. |
| BE_Fla_2018_1 | OK668734 | Belgium | Flandria | Limburg | Maasmechelen | industrial area | 50.995194 | 5.621833 | 2018.07.03. |
| BE_Fla_2018_2 | OK668723 | Belgium | Flandria | Limburg | Maasmechelen | industrial area | 50.995194 | 5.621833 | 2018.07.03. |
| BE_Fla_2018_3 | OK668756 | Belgium | Flandria | Limburg | Maasmechelen | industrial area | 50.995194 | 5.621833 | 2018.07.03. |
| BE_Fla_2018_4 | OK668732 | Belgium | Flandria | Limburg | Maasmechelen | industrial area | 50.995194 | 5.621833 | 2018.07.17. |
| BE_Fla_2018_5 | OK668762 | Belgium | Flandria | Limburg | Maasmechelen | industrial area | 50.995194 | 5.621833 | 2018.07.17. |
| BE_Fla_2018_6 | OK668832 | Belgium | Flandria | Limburg | Maasmechelen | industrial area | 50.995194 | 5.621833 | 2018.07.17. |
| BE_Fla_2018_7 | OK668733 | Belgium | Flandria | Limburg | Maasmechelen | industrial area | 50.995194 | 5.621833 | 2018.06.19. |
| BE_Fla_2018_8 | OK668735 | Belgium | Flandria | Limburg | Maasmechelen | industrial area | 50.995194 | 5.621833 | 2018.06.19. |
| BE_Fla_2018_9 | OK668722 | Belgium | Flandria | Limburg | Maasmechelen | industrial area | 50.995194 | 5.621833 | 2018.06.19. |
| BE_Fla_2018_10 | OK668721 | Belgium | Flandria | Limburg | Maasmechelen | industrial area | 50.995194 | 5.621833 | 2018.06.19. |
| GenBank | JF430393 | Belgium | Flandria | Limburg | Maasmechelen | industrial area | 50.996261 | 5.619860 | 2008.05.27. |

**S2 Table.** Quality features of the complete genome sequence of *Aedes koreicus* achieved by Oxford Nanopore and Illumina sequencing. The completeness of the genome assembly was evaluated with the Benchmarking Universal Single-Copy Orthologs (BUSCO, v 4.1.2) software, using the Diptera lineage dataset. C: complete; S: single; D: duplicated; F: fragment; M: missing.

| **Sequencing platform** | **Assemblies** | **BUSCO analysis numbers** | | | | |
| --- | --- | --- | --- | --- | --- | --- |
|  |  | C | S | D | F | M |
| Nanopore | mosq.contigs.fasta | 1169 | 1136 | 33 | 659 | 1457 |
| Nanopore+Illumina | pilon (1^st^ iteration) | 2384 | 2325 | 59 | 299 | 602 |
| Nanopore+Illumina | pilon (3^rd^ iteration) | 2448 | 2380 | 68 | 270 | 567 |
|  |  | **BUSCO analysis percentages** | | | | |
|  |  | C (%) | S (%) | D (%) | F (%) | M (%) |
| Nanopore | mosq.contigs.fasta | 35.59 | 34.58 | 1.00 | 20.06 | 44.35 |
| Nanopore+Illumina | pilon (1^st^ iteration) | 72.57 | 70.78 | 1.80 | 9.10 | 18.33 |
| Nanopore+Illumina | pilon (3^rd^ iteration) | 74.52 | 72.45 | 2.07 | 8.22 | 17.26 |

**Appendix S1.** Results of Tajima’s D, Fu and Li's Neutrality Tests calculations.

| Sequences | Tajima’s D | P value | Fu and Li's | | P value | |
| --- | --- | --- | --- | --- | --- | --- |
|  |  |  | D test | F test | D test | F test |
| Full dataset | -2,10058 | < 0.05 | -6,26734 | -5,43285 | < 0.02 | < 0.02 |
| Hungary | -1.46929 | P > 0.10 | -2.56505 | -2.59537 | < 0.05 | < 0.05 |
| Belgium | -0,83418 | > 0.10 | -1,63423 | -1,73384 | > 0.10 | > 0.10 |
| Italy | -1.83949 | P < 0.05 | -3.92005 | -3.77266 | < 0.02 | < 0.02 |
| Germany | -1.39905 | > 0.10 | -1.04259 | -1.33169 | > 0.10 | > 0.10 |

**Appendix S2.** Distribution of the 130 investigated *Aedes koreicus* samples among the 31 different haplotypes (Hap#), “Freq” indicates the number of samples belong to the given haplotype, name of the “Sequences” refers to the GenBank Accession Numbers listed in the S1 Table.

[Hap# Freq. Sequences]

[Hap_1: 1	JF430393_1]

[Hap_2: 1	OK668837_1]

[Hap_3: 34	OK668836_1 OK668760_1 OK668751_1 OK668750_1 OK668749_1 OK668759_1 OK668729_1 OK668728_1 OK668727_1 OK668726_1 OK668725_1 OK668724_1 OK668731_1 OK668723_1 OK668722_1 OK668721_1 OK668762_1 OK668755_1 OK668754_1 OK668752_1 OK668761_1 OK668753_1 OK668720_1 OK668719_1 OK668718_1 OK668717_1 OK668716_1 OK668715_1 OK668714_1 OK668713_1 OK668730_1 OK668748_1 OK668825_1 OK668824_1]

[Hap_4: 1	OK668835_1]

[Hap_5: 1	OK668834_1]

[Hap_6: 18	OK668832_1 OK668747_1 OK668745_1 OK668744_1 OK668743_1 OK668741_1 OK668736_1 OK668756_1 OK668733_1 OK668734_1 OK668735_1 OK668732_1 OK668742_1 OK668740_1 OK668739_1 OK668738_1 OK668737_1 OK668746_1]

[Hap_7: 1	OK668831_1]

[Hap_8: 1	OK668830_1]

[Hap_9: 1	OK668828_1]

[Hap_10: 1	OK668819_1]

[Hap_11: 15	OK668818_1 OK668817_1 OK668812_1 OK668811_1 OK668810_1 OK668809_1 OK668774_1 OK668773_1 OK668772_1 OK668771_1 OK668770_1 OK668769_1 OK668768_1 OK668776_1 OK668775_1]

[Hap_12: 30	OK668814_1 OK668797_1 OK668796_1 OK668795_1 OK668791_1 OK668788_1 OK668787_1 OK668786_1 OK668785_1 OK668784_1 OK668783_1 OK668782_1 OK668801_1 OK668800_1 OK668790_1 OK668789_1 OK668781_1 OK668780_1 OK668804_1 OK668779_1 OK668778_1 OK668777_1 OK668798_1 OK668793_1 OK668792_1 OK668802_1 OK668799_1 OK668815_1 OK668803_1 OK668827_1]

[Hap_13: 1	OK668808_1]

[Hap_14: 1	OK668807_1]

[Hap_15: 2	OK668806_1 OK668805_1]

[Hap_16: 6	OK668763_1 OK668766_1 OK668765_1 OK668764_1 OK668767_1 OK668816_1]

[Hap_17: 1	OK668794_1]

[Hap_18: 1	OK668757_1]

[Hap_19: 1	OK668758_1]

[Hap_20: 1	OK668820_1]

[Hap_21: 1	OK668712_1]

[Hap_22: 1	OK668710_1]

[Hap_23: 1	OK668822_1]

[Hap_24: 1	OK668821_1]

[Hap_25: 1	OK668709_1]

[Hap_26: 1	OK668823_1]

[Hap_27: 1	OK668711_1]

[Hap_28: 1	OK668813_1]

[Hap_29: 1	OK668829_1]

[Hap_30: 1	OK668826_1]

[Hap_31: 1	OK668833_1]

**Appendix S3.** Calculated Bayes Factors for specific dispersal routes of *Aedes koreicus* within the investigated European populations.

| **From** | **To** | **Bayes Factor** |
| --- | --- | --- |
| Maasmechelen  (Flandria, Belgium) | Arba  (Friuli Venzia Giulia, Italy) | 31.221086662999646 |
| Pécs  (Southern Transdanubia, Hungary) | Baon  (Veneto, Italy) | 5.9431444460479055 |
| Budapest  (Central-Hungary, Hungary) | Arba  (Friuli Venzia Giulia, Italy) | 7.776197934391781 |
| Budapest  (Central-Hungary, Hungary) | Castel Ivano  (Trentino-Alto Adige, Italy) | 416.67334770003663 |
| Villamontagna  (Trentino-Alto Adige, Italy) | Baone  (Veneto, Italy) | 6.0750831084945105 |
| Maniago  (Friuli Venzia Giulia, Italy) | Baone  (Veneto, Italy) | 46.65826894455457 |
| Pedavena  (Veneto, Italy) | Grigno  (Trentino-Alto Adige, Italy) | 9.250757520355833 |
| San Michele all’Adige  (Trentino-Alto Adige, Italy) | Grigno  (Trentino-Alto Adige, Italy) | 4.673723620251378 |
| San Michele all’Adige  (Trentino-Alto Adige, Italy) | Valli del Pasubio  (Veneto, Italy) | 3.4058369274555003 |
| Grigno  (Trentino-Alto Adige, Italy) | Castel Ivano  (Trentino-Alto Adige, Italy) | 5502.518188580577 |
| Arba  (Friuli Venzia Giulia, Italy) | Castel Ivano  (Trentino-Alto Adige, Italy) | 5.583224432606171 |
| Monteforte d'Alpone  (Veneto, Italy) | Baone  (Veneto, Italy) | 6.300992241289952 |
| Castel Ivano  (Trentino-Alto Adige, Italy) | Valli del Pasubio  (Veneto, Italy) | 4.22011005416697 |
| Castel Ivano  (Trentino-Alto Adige, Italy) | Baone  (Veneto, Italy) | 51.15628021169566 |
| Castel Ivano  (Trentino-Alto Adige, Italy) | Črniče  (Goriška, Slovenia) | 8.022727454894238 |
| Lovrenc na Dravskem polju  (Drava, Slovenia) | Baone  (Veneto, Italy) | 8.78143589286883 |
| Wiesbaden  (Hessen, Germany) | Valli del Pasubio  (Veneto, Italy) | 1829.4080257224512 |
| Wiesbaden  (Hessen, Germany) | Baone  (Veneto, Italy) | 5.8260199803535215 |
| Valli del Pasubio  (Veneto, Italy) | Baone  (Veneto, Italy) | 30.35917468328573 |
| Valli del Pasubio  (Veneto, Italy) | Vittorio Veneto  (Veneto, Italy) | 8.299160285144584 |
